# The CoLoMoTo Interactive Notebook: Accessible and Reproducible Computational Analyses for Qualitative Biological Networks

**DOI:** 10.1101/290411

**Authors:** Aurélien Naldi, Céline Hernandez, Nicolas Levy, Gautier Stoll, Pedro T. Monteiro, Claudine Chaouiya, Tomáš Helikar, Andrei Zinovyev, Laurence Calzone, Sarah Cohen-Boulakia, Denis Thieffry, Loïc Paulevé

## Abstract

Analysing models of biological networks typically relies on workflows in which different software tools with sensitive parameters are chained together, many times with additional manual steps. The accessibility and reproducibility of such workflows is challenging, as publications often overlook analysis details, and because some of these tools may be difficult to install, and/or have a steep learning curve. The CoLoMoTo Interactive Notebook provides a unified environment to edit, execute, share, and reproduce analyses of qualitative models of biological networks. This framework combines the power of different technologies to ensure repeatability and to reduce users’ learning curve of these technologies. The framework is distributed as a Docker image with the tools ready to be run without any installation step besides Docker, and is available on Linux, macOS, and Microsoft Windows. The embedded computational workflows are edited with a Jupyter web interface, enabling the inclusion of textual annotations, along with the explicit code to execute, as well as the visualisation of the results. The resulting notebook files can then be shared and re-executed in the same environment. To date, the CoLoMoTo Interactive Notebook provides access to software tools including GINsim, BioLQM, Pint, MaBoSS, and Cell Collective for the modelling and analysis of Boolean and multi-valued networks. More tools will be included in the future. We developed a Python interface for each of these tools to offer a seamless integration in the Jupyter web interface and ease the chaining of complementary analyses.

## 1 Introduction

Recently, the scientific community has been increasingly concerned about difficulties in reproducing already published results. In the context of preclinical studies, observed difficulties to reproduce important findings have raised controversy (see *e.g.* [7, 15, 40, 43], and [8] for a review on this topic). Although not invalidating the findings, these observations have shaken the community. In 2016, a Nature survey pointed to the multi-factorial origin of this “reproducibility crisis” [4]. Factors related to computational analyses were highlighted, in particular the unavailability of code and methods, along with the technical expertise required to reproduce the computations.

The scientific community is progressively addressing this problem. Prestigious conferences (such as two major conferences from the database community, namely, VLDB^1^ and SIGMOD^2^) and journals such as PNAS^3^, Biostatistics [38], Nature [41] and Science [54], to name only a few, now encourage or even require published results to be accompanied by all the information necessary to reproduce them.

While the reproducibility challenges have first been observed in domains where deluge of data were quickly becoming available (*e.g.*, Next Generation Sequencing data analyses), the problem is now present in many (if not all) communities where computational analyses and simulations are performed. In particular, the System Biology community is facing a proliferation of approaches to perform a large variety of tasks, including the development of dynamical models, complex simulations, and multiple comparisons between varying conditions of model variants. Consequently, reproducing results from systems biology studies becomes increasingly difficult. Furthermore, although the combination of different tools would provide various new scientific opportunities, this is currently hindered by technical issues.

Several initiatives have been launched by the community to address reproducibility issues for computational modelling of biochemical networks. These include guidelines for model annotations (MIRIAM, [27]) and simulation descriptions (MIASE, [51]), as well as standards for model exchange (SBML, [23]) and simulation parametrizations (SED-ML, [52]). This collective effort is coordinated by the *COmputational Modeling in BIology NEtwork* (COMBINE^4^).

The Consortium for Logical Models and Tools (CoLoMoTo^5^) has been organized to bring together computational modelling researchers and address the aforementioned reproducibility and reusability issues within the sub-domain of logical models and software tools [33]. As a first outcome to foster model exchange and software interoperability, the SBML L3 package qual was developed [9, 10]. In this manuscript, we report the next phase of the CoLoMoTo efforts in the area of reproducibility in computational systems biology: The CoLoMoTo Interactive Notebook, which provides an easy-to-use environment to edit, execute, share, and reproduce analyses of qualitative models of biological networks by seamlessly integrating various logical modelling software tools.

The teams involved in CoLoMoTo, gathering around 50 researchers within 20 groups and laboratories, have produced various software tools for the qualitative modelling and analysis of biological networks. They are also involved in the development of novel computational methods and models. This method article presents a collective effort to provide the community with a reproducibility-oriented framework combining software tools related to logical modelling. This framework combines the power of different approaches to ensure repeatability and to reduce the requirement of technical knowledge from users. The provided Docker image facilitates the stability of a contained environment needed for repeatable computational modelling and analyses. The framework includes a set of pre-installed tools from the CoLoMoTo community. On the other hand, specific binding and interfaces integrated in a Jupyter environment reduce the learning curve and improve accessibility. The use of this framework is demonstrated by a case study in a companion protocol article, which consists in a thoroughly annotated Jupiter notebook [28]^6^.

The method article is structured as follows. Section 2 provides a brief introduction to qualitative models of biological networks and to their analyses. Section 3 describes the main components (Docker image, Python programming interface, Jupyter interactive web interface) of our framework to facilitate the access to CoLoMoTo software tools, a prime prerequisite for the reproducibility of the computational analyses. Section 4 illustrates how our framework can address several challenges related to the reproducibility of computational analyses, ranging from the repeat of a sequence of analyses in the exact same software environment, to the use of alternate methods to reproduce a similar result. Finally, Section 5 provides an introductory guide on how to use the new framework, and Section 6 discusses possible extensions.

## 2 Qualitative Dynamical Models of Biological Networks

Since the pioneering work of Kauffman [24], Thomas [49], and others, logical (e.g., Boolean) models have emerged as a framework of choice to model complex biological networks, focusing for example on the roles of transcriptional regulatory circuits in cell differentiation and development, signalling pathways in cell fate decisions, etc. (for a review, see *e.g.* Abou-Jaoudé et al. [2]).

### 2.1 Qualitative modelling

The definition of a qualitative logical model, such as a *Boolean model* usually relies first on the delineation of a *regulatory graph*, where each *node* denotes a regulatory component (*e.g.* a protein or a gene) connected with (positive or negative) *arcs* denoting interactions (activation or inhibition) between its source and target nodes. Each node is modelled as a discrete variable, having a finite number of possible values, typically Boolean, *i.e.*, only two values, 0 or 1, denoting *e.g.* protein absence/inactivity or presence/activity. A *Boolean function* or *rule* is then defined for each node to specify how its value evolves depending on the values of its regulators.

The *state* of a network is modelled as a vector encompassing the (Boolean or multi-valued) values of all the nodes of the regulatory graph, with a prescribed ordering. The state of the network can be updated according to the logical functions defined for each node, triggering a *transition* towards a successor state. When at a given state, several nodes are called for an update, different updating modes can be considered. The *synchronous updating mode* updates all nodes simultaneously, thus leading to a unique successor state. Hence, the dynamical behaviour is fully deterministic. In contrast, the *asynchronous updating mode* updates only one node, choosen non-deterministically, thus leading to different possible successor states. Several variants and extensions of these updating mode have been defined, for instance assigning pre-determined priorities or assigning probabilities to nodes updates, or considering simlultaneous updates of sub-groups of nodes.

### 2.2 Dynamical analysis

The dynamical behaviour of the model can be represented by its *state transition graph*, where vertices are the different states of the network, and directed edges are the possible *transitions* between states, following a selected updating mode. *Dynamical analyses* consist then in characterizing different properties of this state transition graph.

*Attractors* are one of the most prominent features studied in Boolean and multi-valued networks. Attractors model the asymptotic behaviours of the system, and correspond to the *terminal strongly connected components* of the state transition graph. Attractors can be of different nature, either reduced to a single *stable state* (or *fixpoint*), from which no transition are possible, or *cyclic* sequences of states, modelling sustained oscillations. From a biological point of view, computing attractors is generally particularly relevant. The presence of multiple (disjoint) attractors can represent alternative cell fates (such as cell differentiation states), while cyclic attractors further represent periodic behaviours (such as cell cycle or circadian rhythms). The computation of attractors is addressed by different software tools, such as bioLQM [31], GINsim [32], Pint [36], BoolSim [19], BooleanNet [3], pyBoolNet [25], and BoolNet [30].

*Simulations* allow capturing the states which are *reachable* from a given (set of) *initial state(s)*. They can consist of random walks in the complete state transition graph and take into account updating priority schemes to distinguish fast versus slow processes and thereby obtain a simpler state transition graph [16] as implemented in the software tool bioLQM, or user-defined transition probabilities and timing, as implemented into the software tools MaBoSS [46, 47] and CellCollective [22, 50].

*Model checking* techniques developped for software verification in computer science allow verifying formally dynamical properties on state transition graphs and are regularly employed for analysing biological systems [1, 5, 6, 13]. The properties are specified using so-called *temporal logics* which enable the formulation of queries regarding asymptotic or transient dynamical properties and taking into account all the state transitions of the model. The accordance of a Boolean/multi-valued model with such properties is verified using a general purpose model checker such as NuSMV [11] for which GINsim and Pint provide access to.

It is worth noticing that the number of states of a Boolean or multi-valued network grows exponentially with the number of nodes. The above mentioned methods typically suffer from this complexity, and can be limited with the size of the networks (currently, mostly around fifty to a hundred of nodes, depending on the analysis and the complexity of the dynamics).

Nevertheless, different approaches enable the analysis of large scale qualitative networks, for instance by using *model reductions*, such as by Naldi et al. [34] which preserves stable states, while cyclic attractors and reachability can be affected in predictable ways, as implemented in bioLQM, or by using *formal approximations* of dynamics, as implemented in Pint, which allow tackling networks with several thousands of nodes [36, 37].

Figure 1 gives an overview of a range of software tools for analysis of qualitative models, specifying their main features along with the main underlying technologies.

**Figure 1:**
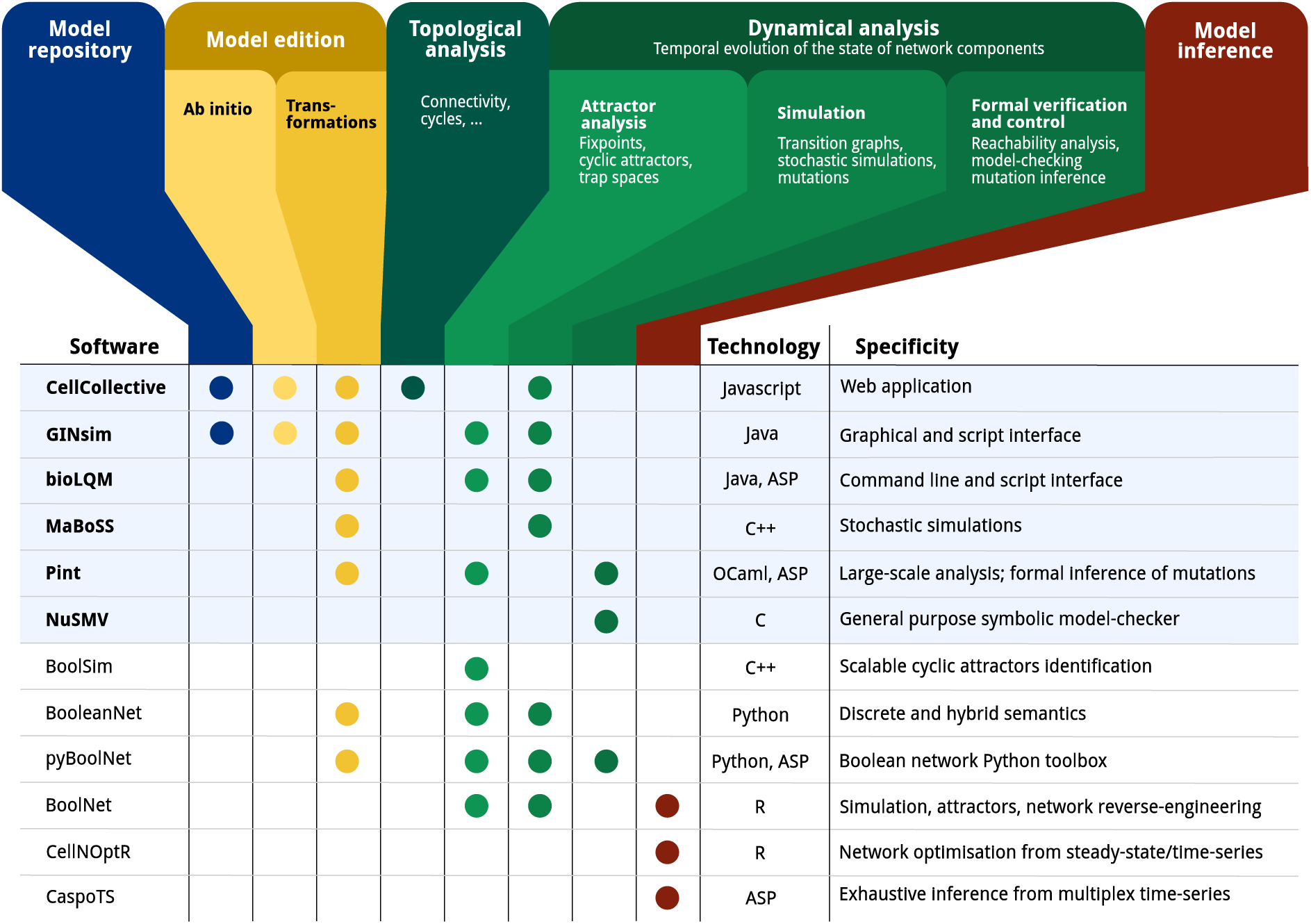
Feature matrix and specifities of a range of software tools related to qualitative biological networks: CellCollective [22]; GINsim [32]; bioLQM [31]; MaBoSS [46]; Pint [36]; NuSMV [11]; BoolSim [19]; BooleanNet [3]; pyBoolNet [25]; BoolNet [30]; CellNOpt [48]; CaspoTS [35]. The current CoLoMoTo Docker (2018-03-31) ships the software indicated with a bold font and a light blue background. “Model repository” refers to searchable databases of models; “Model edition” refers to the features related to creating and modifying a qualitative model, where “*ab initio”* refers to the interactive model building from scratch, and “transformations” refers to operations such as mutations, Booleanization, model reduction, etc. “Topological analysis” refers to the extraction of features from the regulatory graph, such as the different feedback cycles, graph theory measures, etc. “Dynamical analysis” refers to properties related to the state transition graph of Boolean/multi-valued networks, where “Attractor analysis” refers to the identification of stable states, cyclic attractors, and related features; “Simulation” refers to the sampling of trajectories within the state transition graph, possibly parameterized with stochastic rates and mutations; “Formal verification and control” refers to exhaustive analyses for assessing strictly temporal properties, such as reachability, and deducing mutations for controlling the system. Finally, “Model inference” refers to the derivation of Boolean/multi-valued network which are compatible with given properties and observation data.

## 3 Accessibility of CoLoMoTo software tools

The CoLoMoTo Interactive Notebook aims at offering a unified environment for accessing a range of complementary software tools related to the analysis of qualitative models of biological networks. To achieve such a goal, our framework relies on three complementary technologies.

First, we use the Docker system to provide images of pre-installed selected CoLoMoTo software tools, thus reducing significantly the burden of installing individually each software tool. The software installed within Docker images can be executed on GNU/Linux, macOS, and Microsoft Windows, and can be accessed by standard workflow systems, such as SnakeMake [26].

Then, we developed a collection of Python modules to provide a unified interface to the features of the selected software tools. The Python modules allow to parametrize and execute the different analyses, and fetch their results which can be then further processed, including by a different tool through its respective Python module.

This uniform Python interface is particularly relevant in the Jupyter web interface [39], where it allows editing executable notebooks on qualitative biological networks by seamlessly combining different software tools.

### 3.1 The CoLoMoTo Docker image

Overall, we witness a growing ecosystem of software tools based on different technologies and offering a wide range of complementary features. Noteworthy, these tools typically rely on tailored formalisms and settings, which enable specific methods but at the same time affect the results. One obvious example is the consideration of a specific updating mode, as synchronous and asynchronous dynamics may differ extensively. Furthermore, to address increasingly large networks, many tools rely on advanced data-structures and resolution methods, which are implemented in dedicated software libraries. The distribution of these tools then become quite challenging, as they rely on numerous dependencies, some of which are difficult to install or available only for a few operating systems (most of the time GNU/Linux).

The Docker container technology allows to circumvent such distribution issues by providing a mean to supply pre-installed and fully configured software environments in so-called *Docker images*. On GNU/Linux, the execution of a Docker image consists mainly in executing the software in an isolated environment, requiring no operating system virtualization. Therefore, the overhead of using Docker on GNU/Linux is close to zero. A Docker image can also be executed on macOS or Microsoft Windows without any modification. On these operating systems, Docker relies on virtualization technologies, which are relatatively lightweight and result in limited performance loss on recent hardware.

The current CoLoMoTo Docker image colomoto/colomoto-docker:2018-03-31 contains the follow-ing pre-installed software for the logical modelling and analsyis of biological networks: GINsim [32]; bioLQM [31]; CellCollective [22]; MaBoSS [46]; Pint [36]; and NuSMV [11].

The CoLoMoTo Docker image then provides access to these tools without requiring any installation step beside installing Docker^7^. For instance, the Docker image can be used in association with a workflow manager to chain and run a series of software functionalities. Supplementary file “*SnakeMake”* provides an example of SnakeMake workflow relying on GINsim and NuSMV in the CoLoMoTo Docker.

An important challenge is the maintenance and extendibility of such Docker images to reduce the complexity of upgrading or adding software tools with their respective dependencies. To that aim, we require that each software tool is independently packaged for GNU/Linux using the *Conda package manager*^8^. We then rely on the dependency management system of Conda to ensure that the correct pre-requisites are installed in the Docker image^9^. A beneficial side effect of this technical choice is that the aforementioned software tools can be installed on GNU/Linux platforms using Conda, without using Docker.

### 3.2 A unified interface for calling and chaining tools with Python

The software tools considered for the CoLoMoTo Docker image present different interfaces: CellCollective is a web application, GINsim has a graphical user interface along with a scripting interface, bioLQM has a command line and a scripting interface, Pint has a command line and a Python interface, MaBoSS has a command line interface.

GINsim, CellCollective and bioLQM support the SBML-qual format, while bioLQM provides the conversion of a standard SBML-qual model into Pint or MaBoSS model formats, thereby enabling the exchange of models between all these tools.

The recourse to different interfaces complicates the design of a model analysis combining multiple tools. To address this issue, we have developed a Python interface for each of the tool embedded in the CoLoMoTo Docker image, which greatly ease the execution of different tool functionalities, fetch the results, and use these later as input for other executions.

Each tool comes with a dedicated Python module, providing a set of functions to invoke the underlying software tool appropriately. Therefore, from a single Python shell, one can invoke and chain analyses performed by different tools. This can be seen as an improved command line interface, greatly enhanced by the use of intermediate Python objects. Such an approach also promotes the use of standard Python data-structures to store objects such as model states or graphs, which can then be processed by common Python libraries, *e.g.*, Pandas^10^ or NetworkX^11^.

Hereafter, we give an overview of the resulting Python programming interface, focusing on the general model input mechanism and the main features implemented for each of the software tools.

#### Model input and tool conversions

Despite their very different features, all the tools considered here take as input a logical model, in an adequate format. All the related Python modules provide a load function, which takes as input the location of the model, being a local file, for instance:

~~~
m = biolqm.load(“path/to/localfile.sbml”)
~~~

a web link to a file, as obtained on GINsim repository^12^ for instance:

~~~
m = biolqm.load(“http://ginsim.org/sites/default/…MamCC_Apr2016.sbml”)
~~~

or a web link to the model on CellCollective, for instance:

~~~
m = biolqm.load(“https://cellcollective.org/#5128/lac-operon”)
~~~

In each case, the returned object (identified by m in the above examples) is a Python object representing the loaded model and defined specifically for the corresponding tool (Python module). Table 1 lists the supported input format for each software tool.

**Table 1:** Model input formats for the software tools included in the CoLoMoTo Docker image

| Software tool | Supported input formats |
| --- | --- |
| BIOLQM | SBML-qual (.sbml), raw logical functions, truth table |
| GINSIM | GINML (.ginml, .zginml) |
| PINT | Automata network (.an) |
| MABOSS | Dedicated network/configuration files (.bnd/.cfg) |
| NUSMV | SMV file (.smv) |

When possible, Python modules provide functions to convert a model for a compatible tool. These functions are of the form moduleA.to moduleB(modelA). Figure 2 lists the currently supported model conversions. The following Python code shows an example of usage:

**Figure 2:**
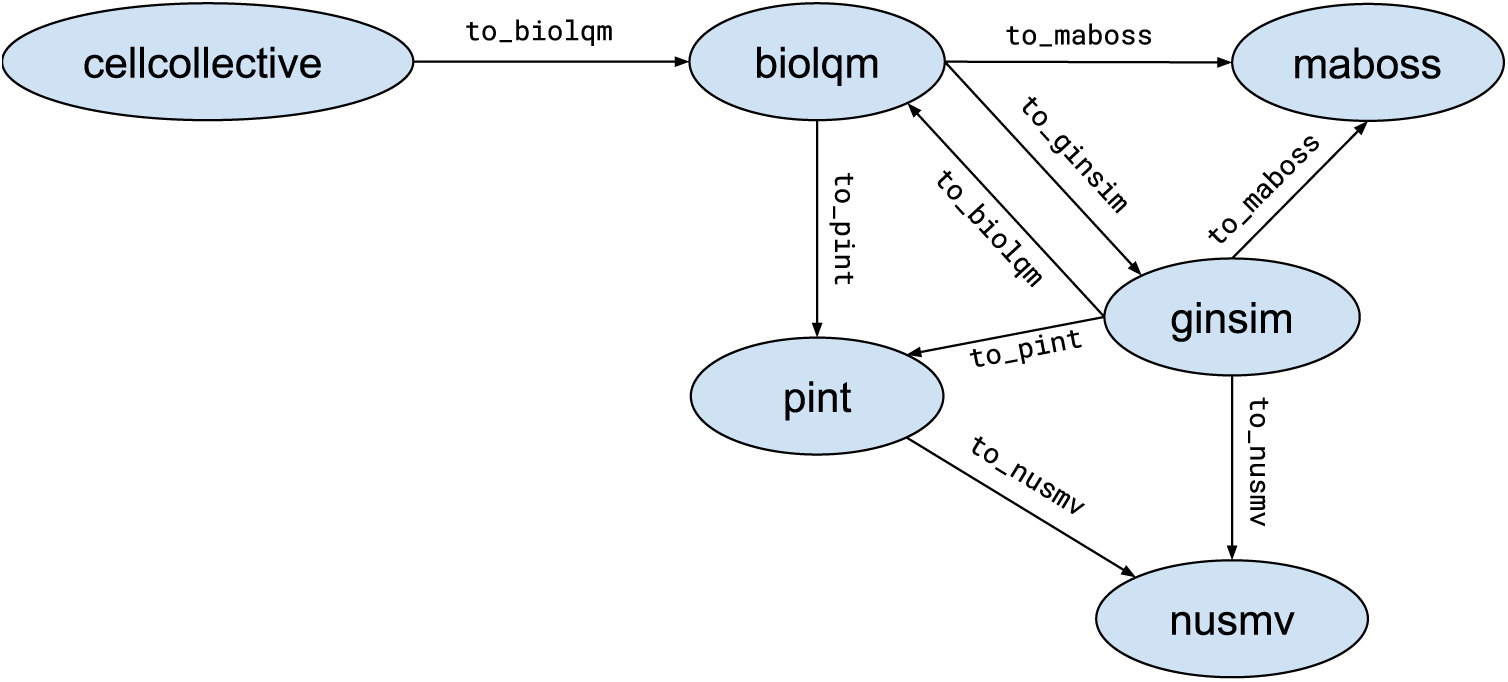
Supported conversion between Python module models

~~~
lrg = ginsim.load(“http://ginsim.org/sites/default/…MamCC_Apr2016.zginml”)
lqm = ginsim.to_biolqm(lrg)
an = biolqm.to_pint(lqm)
~~~

Here, lrg is a Python object representing a GINsim model, lqm is a Python object representing a bioLQM model, and an is a Python object representing a Pint model.

#### CellCollective– Modelling platform, repository, and knowledge base

The cellcollective Python module allows connecting to the CellCollective [22] web application (https://www.cellcollective.org), in order to download the models in SBML-qual format and extract network nodes meta-data (*e.g.*, UnitProt identifiers) when available.

The supplementary file “*Notebooks/demo-cellcollective”*^13^ provides a brief demonstration of the Python module usage.

#### GINsim– Regulatory network modelling

The ginsim Python module provides direct access to the Java programming interface of GINsim [32]. GINsim is available and documented at http://www.ginsim.org. In particular, besides the export of a GINsim model into various file formats, the Python module allows to visualise the network regulatory graph, with the activation and inhibition relationships between the nodes. The visualisation function (ginsim.show) optionally takes as argument a Python dictionary associating a level with each node; then, the nodes of the network are coloured according to these levels.

This is illustrated in the supplementary file “*Notebooks/demo-ginsim”*^14^.

#### bioLQM– Qualitative model toolbox

The biolqm Python module provides direct access to the Java programming interface of bioLQM [31]. bioLQM is available and documented at http://colomoto.org/biolqm. bioLQM supports the conversion of SBML-qual files, GINML files, as well as simple textual files specifying the raw logical functions into the formats associated with the different software tools.

Besides the file format features, bioLQM implements *model modifications*, such as mutations forcing the value of given nodes, iterative model reduction(see above), model reversal, the conversion of multivalued model into Boolean ones, as well as the computation of *stable states*, *trap spaces*, and *simulations*.

Part of these features are illustrated in the supplementary file “*Notebooks/demo-bioLQM”*^15^.

#### Pint– Formal predictions for controlling trajectories

`The pypint` Python module provides complete access to Pint [36] features documented at https://loicpauleve.name/pint. The software Pint is devoted to the analysis of trajectories in very large-scale asynchronous Boolean and multi-valued networks. Its main features include the verification of the existence of a trajectory reaching a state of interest (*reachability*), the identification of common points between *all* the trajectories leading to a state of interest (*cut sets*), and the *formal prediction of mutations* preventing the existence of any trajectory to the given state.

These features are illustrated in the supplementary file “*Notebooks/demo-pint”*^16^.

#### NuSMV– Model verification

The nusmv Python module provides a simple interface to the NuSMV model checker [11] for verifying LTL (trace) and CTL (computation tree) temporal logic properties. The specification of LTL and CTL properties can be facilitated using the colomoto.temporal logics Python module, which takes advantage of Python objects for the different logical operators and ease their combination.

Let us consider the following example using the CTL operators from the aforementioned Python module:

~~~
p1 = AG (S(a=1))
p2 = EF (p1)
~~~

Here, the variable p1 is a CTL formula specifying that the node *a* is active (S(a=1)) in all the reachable states (AG operator). The variable p2 is a CTL formula specifying that there exists a trajectory leading to a state (EF operator) from which the property p1 is verified.

In the above example, S specifies a property on a state, by giving the values of some nodes of the network. The conversion of a network model into NuSMV format depends on the tool used, sometimes introducing different variable names for the nodes of the original biological network. But this technical point is transparent for the user: the nusmv Python module will automatically translate the node names into the correct NuSMV variable names.

The supplementary file “*Notebooks/demo-nusmv”*^17^ gives a simple example of usage of the nusmv

Python module to verify properties of a GINsim model.

#### MaBoSS– Stochastic simulations

The maboss Python module provides an interface to MaBoSS [46], available at https://maboss.curie.fr, as well as basic plotting functionalities. The purpose of MaBoSS is to perform stochastic simulations of a Boolean network where the propensity of transitions (probabilistic rates) are explicitly specified. The Python module allows to fully define and parameterize a model, as well as parsing an existing MaBoSS model and modify it programmatically. The object returned after the simulations can then be used to plot the probability of state activation over time, and the proportion of states in which the simulations ended, in order to estimate the probability of reaching different attractors.

The supplementary file “*Notebooks/demo-maboss”*^18^ provides a brief tutorial to the main features of the maboss Python module.

#### Advanced combinations of tools

These Python modules provide a unified interface to chain different tools and process their results. The small tutorials referenced above show simple chaining of tools, most of the time using a tool to import a model (e.g. from CellCollective or GINsim) and convert it (using bioLQM) for specific analysis by another tool.

As the Python functions of the different modules relies on standard Python data-structures, such as list and dictionaries, it is possible to easily re-use the result from a tool function as input to the function of a different tool. A simple example is provided in supplementary file “*Notebooks/demo-ginsim”*^19^, where we use bioLQM to compute the fixpoints of a GINsim model, and then give one of the resulting state as input to GINsim show function to display the computed fixpoint in the regulatory graph.

Moreover, one can use the programmatic features of Python to implement advanced algorithms for executing multiple analyses and process their result. For instance, one can program loops to iterate over a list of results of a preceding analysis from one tool to perform a subsequent analysis on each result with another tool. This is illustrated in the supplementary file “*Notebooks/demo-pint+maboss”*^20^ where we use Pint to formally predict combinations of mutations controlling the existence of trajectories towards a specified state; then, we quantify with MaBoSS the efficiency of applying only partially the predicted combinations, by evaluating each related double-mutants. The example involves Python for loops and a function to enumerate all possible subsets provided by the standard Python library. The notebook also relies on CellCollective to fetch the model, and on bioLQM to perform the adequate model conversions.

### CoLoMoTo Jupyter interactive notebook

*Jupyter*^21^ is a software providing an interactive web interface for creating documents, called *notebooks*, mixing code, equations, and formatted texts. A notebook typically describes a full analysis workflow, combining textual explanations, the code itself, along with parameters to reproduce the results. A notebook is a single file that can be easily modified, shared, re-executed and visualised online. The short tutorials of the previous section provided in the supplementary file “*Notebooks”* are actually Jupyter notebooks (files with the extension `.ipynb`) and can be re-executed using Jupyter.

A Jupyter notebook is sequence of so-called *cells*, which can contain formatted text, including sections, links, images, tables, etc., or which can contain code in a specified programming language, typically Python. A code cell can be executed (by pressing Shift-Enter) and the value returned by the code is displayed below the cell. The display format is selected according to the type of the returned value (image, graph, list, table, #x2026;) to offer an adequate visualisation.

Having a unified Python interface to invoke the CoLoMoTo software tools, one can directly create Jupyter notebooks for the analysis of qualitative biological networks using these tools, as shown in supplementary file “*Notebooks”* and in the companion publication [28] providing a complete model analysis workflow.

To improve the user experience for editing Jupyter notebooks, we added several features in the CoLoMoTo Python modules, which increase interactivity. First, menus provide pre-defined Python code for accessing to the main features of the tools. Figure 3 shows a screenshot during the edition of a Jupyter notebook with its graphical interface. Next, we added the possibility to interactively upload a model file. This feature is particularly useful when used in combination with Docker or on a remote server with no direct access to the user file system. Finally, some Python modules, in particular maboss Python model, provides JavaScript widgets to generate Python code interactively.

**Figure 3:**
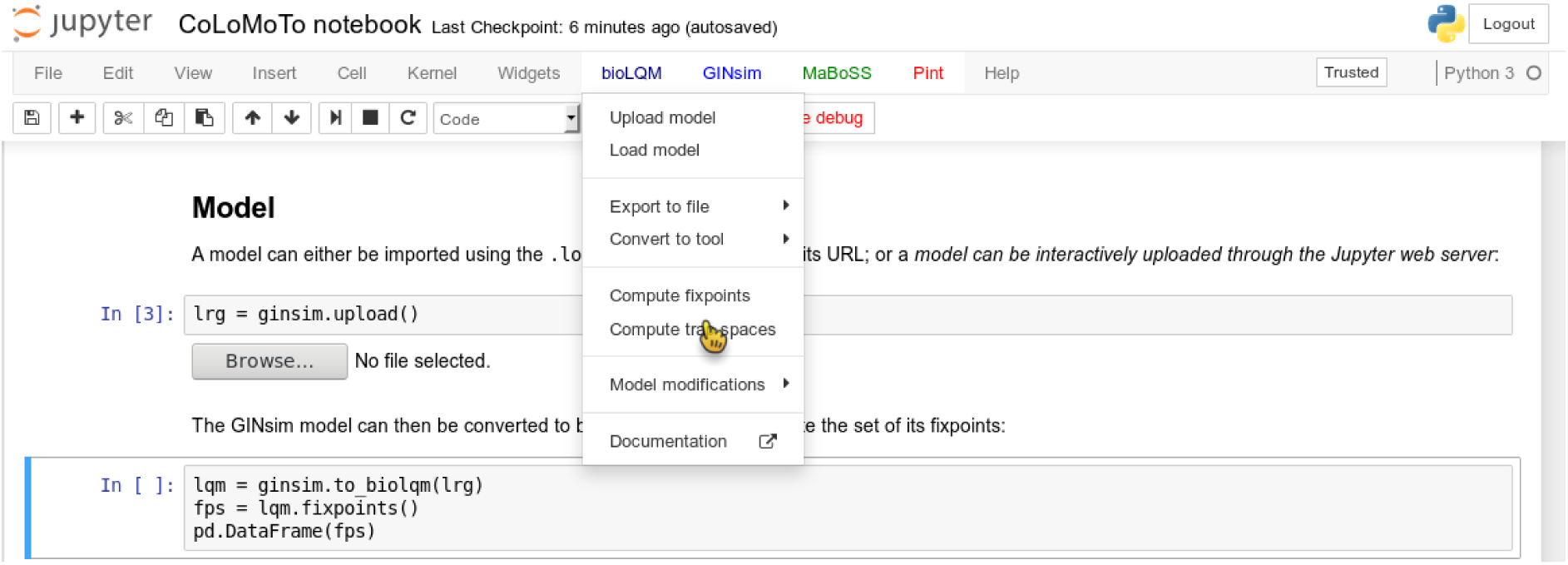
Screenshot during a Jupyter notebook edition showing the menu of the bioLQM tool

The Jupyter notebook server is included in the CoLoMoTo Docker image (see the Discussion section for a quick usage guide), while a public demonstration web instance is available at http://tmpnb.colomoto.org.

### 4 Reproducibility of computational analyses

#### 4.1 From repeatability to reproducibility

The literature provides a range of definitions for the reproducibility of *in silico* experiments by analogy to wet lab experiments [12, 14, 17, 18, 21, 29, 44]. Four *levels* of reproducibility are then commonly distinguished.

An *in silico* experiment is said to be **repeated** when it is performed using the same computational set-up as the original experiment. The major goal of the *repeat* task is to check whether the initial experimental result was correct and can be obtained again. The difficulty lies in recording as much information as possible to repeat the experiment so that the same conclusion can be drawn. Interestingly, [17] discusses the granularity at which information (experiments, data sets, parameters, environment) should or could be recorded and underlines the fact that the key point is to determine the right balance between the effort required to record information and the capability of obtaining identical results.

An *in silico* experiment is said to be *replicated* when it is performed in a new setting and computational environment, although similar to the original ones. When it can be successfully replicated, a result has a high level of robustness: it remained valid when using a similar (although different) protocol. A continuum of situations can be considered between repeated and replicated experiments.

A result is then defined as ***reproduced***, in the broadest possible sense of the term by denoting the situation where an experiment is performed within a different environment, with the aim to validate the same scientific hypothesis. In other words, what matters here is the conclusion obtained and not the methodology considered reaching it. Completely different approaches can be designed, different data sets can be used, as long as the experiments support the same scientific conclusion. A reproducible result is thus a high-quality result, confirmed in various ways.

A last very important concept related to reproducibility is that of ***reuse***, which denotes the case where a *different* experiment is performed, with similarities with an original experiment. A specific kind of reuse occurs when a single experiment is reused in a new context (and thus adapted to new needs), the experiment is then said to be *repurposed*.

It is worth noticing that *repeating* and *replicating* may appear to be technical challenges compared to *reproducing* and *reusing*, which are the most important scientific objectives. However, before investigating alternative ways of obtaining a result (to reach reproducibility) or before reusing a given methodology in a new context (to reach reuse), the original experiment has to be carefully tested especially by reviewers or any peer), demonstrating its ability to be at least repeated and hopefully replicated [17, 45].

#### 4.2 Repeat analysis in the same software environment

Ensuring that a sequence of computational analyses can be repeated by other scientists several months or years after its publication is difficult. Indeed, besides software availability, the version of the tools can be crucial: a new version of a tool can change the default parameters and even some features so that the published instructions become obsolete.

Whereas a Docker image addresses efficiently the issue of making software available, providing a safe way for repeating a notebook content years after its creation requires additional technical procedures.

First, CoLoMoTo Docker images are constructed by specifying explicitly the version of each software. Furthermore, an automatic validation procedure is performed by checking that a set of notebooks still execute without error. Once validated, the Docker image is then tagged with a time-stamp, typically the date of the image validation (of the form `YYYY-MM-DD`, *e.g.*, 2018-03-31). These tagged images are then stored in the public Docker image registry, and can be retrieved any time later. The list of existing tags of colomoto/colomoto-docker Docker images can be viewed at https://hub.docker.com/r/colomoto/colomoto-docker/tags/.

When sharing a notebook, and notably when attaching it to a publication, it is highly recommended specifying the time-stamp of the Docker image in which the notebook has been executed. Then, by downloading the image with this specific tag, it is ensured that other users will repeat the execution in exactly the same software environment.

To help to follow this recommendation, we took two technical decisions. First, we do not use the default non-persistent tag for Docker images (*latest*). It means that the user has to always specify explicitly the time-stamp of the CoLoMoTo Docker image. To remove the burden of actively checking the list of existing time-stamps, we provide a script which, by default, will fetch the most recent Docker image (see Section 5). Second, when loading a CoLoMoTo-related Python module within a Docker container, a textual message indicating the time-stamp of the Docker image is displayed. Therefore, when created within a CoLoMoTo Docker image, notebooks always contain the required information to repeat their execution.

Because a Jupyter notebook is a single file containing everything to execute it, one can easily check if it can be *replicated* in a different software environment, *i.e.*, using a more recent CoLoMoTo Docker image.

Moreover, a notebook can be easily *repurposed* by modifying some arguments of the Python function calls, for instance changing the input model or analysis parameters. One can even define interactive notebooks describing a common model analysis, so that the user only need to provide the input model and execute the Jupyter code cells, as shown in the Supplementary file “*Notebooks/demo-interactive-fixpoints”*^22^ for the computation and visualisation of the stable states of a bioLQM model.

#### 4.3 Reproduce analysis with a different method

Reproducing a same analysis with two different methods is a good mean to increase confidence in the results, as it reduces the chance of software misuse or that the results are affected by a software bug.

The subset of software tools selected for this first CoLoMoTo Docker image presentation already provides redundant implementations of equivalent model analyses, in particular for the identification of stable states and for the verification of temporal properties with NuSMV. To help switch between two tools for performing the same task, we harmonised the usage of Python module functions to ensure that the same functions with the same arguments generate equivalent results with different tools.

##### 4.3.1 Stable states

There exists several methods to compute the full set of stable states (or fixed points) of a logical model, relying on different data-structures and different algorithms.

The software biolqm implements the computation of stable states for Boolean and multi-value logical model using a Java implementation of decision diagrams. In contrast, the software Pint implements the computation of stable states of Automata networks (a generalisation of logical networks) using Boolean satisfaction constraints. As bioLQM provides a conversion of Boolean/multi-valued network into equivalent Automata networks, it is possible to compute the stable states of a model with both software tools.

Both `bioLQM` and `pypint` Python modules provide a `fixpoint` function which take as input the model instance of the corresponding tool, and both returns a list of Python dictionaries describing the states being fixpoints. Provided lqm is a bioLQM model, the following Python code compute its stable states with both tools:

~~~
fps_biolqm = bioLQM.fixpoints(lqm)
fps_pint = pypint.fixpoints(biolqm.to_pint(lqm))
~~~

The supplementary file “*Notebooks/demo-reproducibility-fixpoints”*^23^ shows a complete example of reproduction of stable state computation using bioLQM and Pint.

##### 4.3.2 Temporal property verification (model-checking)

Both GINsim and Pint allow to export their respective model into NuSMV format, where temporal properties can be specified using LTL or CTL (see Section 2.2). However, as the input formalism for these tool relies on different paradigms (logical rule-centred specification in the case of Boolean/multi-valued networks in GINsim; transition-centred specification (à la Petri nets) in the case of Automata networks in Pint), the generated NuSMV models have different features. Nevertheless, in the appropriate settings, the verification of an equivalent CTL or LTL property should give the same result.

Hence, the functions ginsim.to nusmv and pypint.to nusmv are implemented in such ways that, when using their default options, the resulting NuSMV models, albeit different, should produce identical results for identical temporal logic properties. Note, however, that each tool provides specific options for the NuSMV export, which can lead to incomparable results.

The following Python code uses operators defined in the Python module colomoto.temporal logics to specify a property p meaning that from any state, there always exists a trajectory leading to a cyclic attractor where the level of node *a* can always oscillate. Then, assuming lrg is a GINsim model, the code uses GINsim and Pint conversions to NuSMV to perform model verification.

~~~
p = EF (AG (EF (S(a=0)) & EF (S(a=1))))
~~~

~~~
nusmv_ginsim = ginsim.to_nusmv(lrg)
nusmv_ginsim.add_ctl(p)
nusmv_ginsim.verify()
~~~

~~~
nusmv_pint = pypint.to_nusmv(ginsim.to_pint(lrg))
nusmv_pint.add_ctl(p)
nusmv_pint.verify()
~~~

Note that the Python object p represents the CTL property to be tested, whatever the origin of the model (GINsim or Pint).

The supplementary file “*Notebooks/demo-reproducibility-modelchecking”*^24^ provides a more detailed example of the reproduction of model-checking results using GINsim and Pint.

### 5 Quick-usage guide

On GNU/Linux, macOS, or Microsoft Windows, provided that Docker and Python are installed, a helper script to run the CoLoMoTo Docker image and the embedded Jupyter notebook can be installed and upgraded from a terminal using the following command^25^:

~~~
pip install -U colomoto-docker
~~~

The Docker image and the Jupyter notebook interface can be started by executing the following command in a terminal^26^:

~~~
colomoto-docker
~~~

Without any argument, the command will use the most recent CoLoMoTo Docker image. To use the image at a specific tag, append the -V option (*e.g.*, `colomoto-docker -V 2018-03-31`).

The execution of this command will open a web page with the Jupyter notebook interface, enabling loading and execution of notebooks. A new notebook can be created by using the “New/Python3” menu. In this environment, the user has access to all CoLoMoTo Python modules. A code cell is executed by typing “Shift+Enter”. The menu and toolbar allow quick access to main Jupyter functionalities.

Warning: by default, the files within the Docker container are isolated from the running host computer, and are deleted when stopping the container. To have access to the files of the current directory of the host computer, the option `--bind` can be used:

~~~
colomoto-docker --bind
~~~

The container can later be stopped by pressing Ctrl+C keys in the terminal. See colomoto-docker--help for other options. Additional documentation for running the CoLoMoto Docker image can be found at https://github.com/colomoto/colomoto-docker.

## 6 Discussion

### Academic use cases

The prime aim of the CoLoMoTo Interactive Notebook is to assist with the production of accessible and reproducible computational analysis of biological models, with a focus on qualitative models, including Boolean and multi-valued networks. As demonstrated in supplementary file “*SnakeMake”*, the CoLoMoTo Docker image can also be used in standard workflow systems, such as SnakeMake, to lighten the burden of installing the different software tools and make them accessible on different operating systems.

A notebook issued from the CoLoMoTo Docker image gives some guarantees of repeatibility, as it will contain references to the persistent Docker image to re-execute it in the same software environment. Therefore, the notebook file (with `.ipynb` extension) can be distributed as a supplementary file of the related scientific article with instruction to run the Docker image. The Jupyter interface allows to export the notebook in a static HTML file, which could also be joined to the supplementary file to provide a quick way to visualize it.

A notebook can also be distributed independently, for instance by publishing it on *Gist*^27^ or *myExperiment*^28^ [20], which allow to visualise download, and follow potential updates of the notebook. For instance, the tutorial notebook presented by Levy et al. [28] is hosted at https://gist.github.com/pauleve/a86717b0ae8750440dd589f778db428f. Services like Zenodo^29^ also allow creating persistent DOI links to a specific version of a notebook file.

The CoLoMoTo Interactive Notebook is also relevant for teaching purposes. With Jupyter, students can straightforwardly execute, modify, and extend a template notebook to learn methods for analysing models of biological networks. Docker is a standard technology very often supported by local cloud infrastructures, which can therefore provide dedicated resources to execute remotely and privately the CoLoMoTo Jupyter web interface.

### Extending the CoLoMoTo Interactive Notebook

The CoLoMoTo Docker image can be easily extended to include additional tools. The Docker architecture allows inheriting from an existing container, adding a new layer with additional executables. Contributions are welcome through GitHub^30^. The software tool must be usable from the Jupyter interface and should be able to connect with at least one other tool already included. Furthermore, a demonstration notebook should be provided to illustrate the tool usage and how it can be combined with other tools.

Currently, all the embedded tools require an already defined model. Nevertheless, once loaded, a model can be subsequently modified from the Python interface (see tool feature matrix in Figure 1). We are currently considering the development of a programmatic interface for model definition *ab initio*. One of the main challenge is to provide a decent visualisation of the programmatically-created model.

A potential direction is to include a visual edition module in the Jupyter interface, which represents a substantial development effort.

The support for standard exchange formats is key to enable reproducibility of analyses with different tools. In that sense, bioLQM plays an important role for the CoLoMoTo Interactive Notebook as it provides bridges between SBML-qual standard specifications and numerous software tools (Figure 2). The Tellurium Notebook system by [42] offers support for SED-ML to help reproduce quantitative simulation of biological networks. Future work should consider bringing this feature for qualitative models as well, in order to better meet FAIR (Findability, Accessibility, Interoperability, and Reusability) recommendations [53].

## Author Contributions

AN, CH, DT and LP designed the main principles of the CoLoMoTo Interactive notebook and its distribution. AN, CH, NL and LP implemented the necessary Python modules, their integration in the Jupyter interface, and the Docker image. AN and LP edited the notebook tutorials, while CH edited the SnakeMake workflow example. All authors contribute to the writing of the article under the supervision of DT and LP. All authors reviewed the content of this article and agreed to endorse it.

## Acknowledgments

The authors would like to thank the attendees of the fourth CoLoMoTo meeting in Paris, July 17-19, for the insightful discussions which led to designing the CoLoMoTo Interactive notebook.

## Supplemental Data

The supplemental data “*SnakeMake”* contains an example of SnakeMake workflow which uses the CoLoMoTo Docker image to execute the different analyses.

The supplemental data “*Notebooks”* contains several short Jupyter notebooks which demonstrate different usage of the CoLoMoTo interactive notebook, listed in Table 2. The `.ipynb` files can be imported and executed within the Jupyter interface of the CoLoMoTo notebook, using the Docker image colomoto/colomoto-docker:2018-03-31. For each notebook file is included a static HTML file to quickly preview the Jupyter rendering of the notebook, without any requirement. These notebooks can also be previewed and downloaded at https://nbviewer.jupyter.org/github/colomoto/colomoto-docker/tree/2018-03-31/tutorials.

**Table 2:**
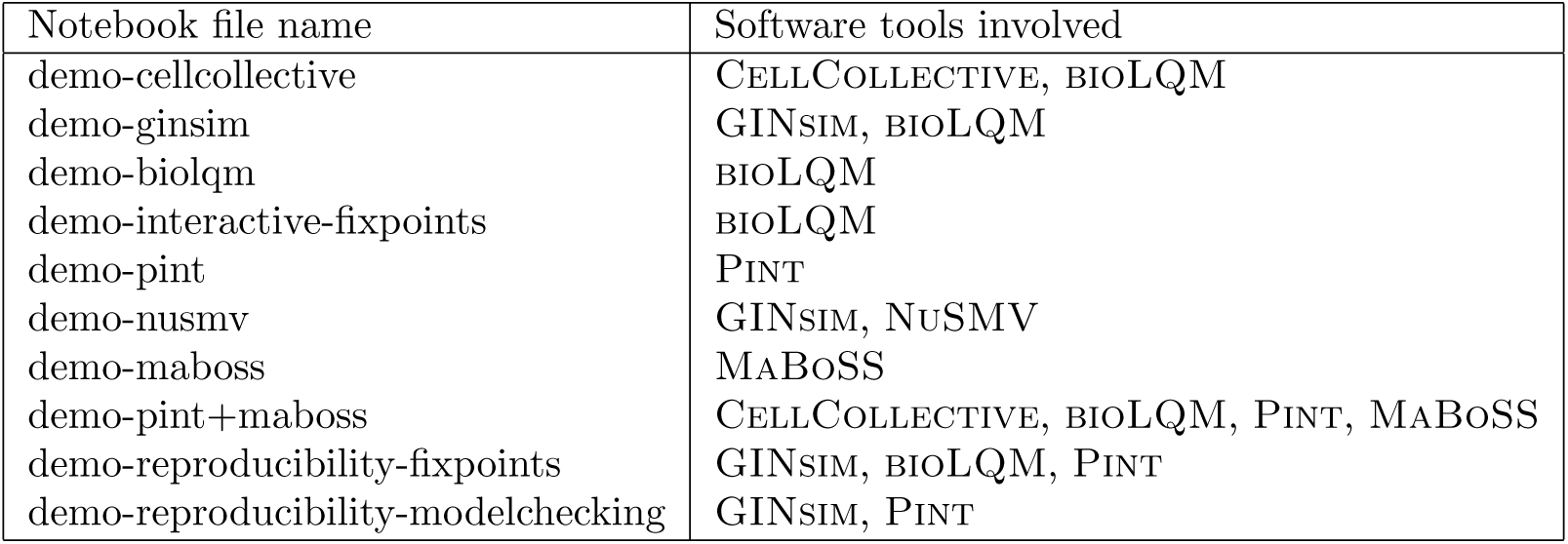
List of notebook files in supplemental data “*Notebooks”* demonstrating some features of the CoLoMoTo Interactive Notebook

## Footnotes

1 International conference on Very Large Data Bases

2 ACM’s Special Interest Group on Management Of Data.

3 http://www.pnas.org/site/authors/format.xhtml

4 http://co.mbine.org

5 http://colomoto.org

6 The notebook can be previewed and downloaded at https://nbviewer.jupyter.org/github/colomoto/colomoto-docker/blob/2018-03-31/usecases/Usecase%20-%20Mutations%20enabling%20tumour%20invasion.ipynb

7 See https://docker.com for installation instructions

8 https://conda.io

9 CoLoMoTo-related conda packages are available in the colomoto conda channel. See https://anaconda.org/colomoto

10 https://pandas.pydata.org/

11 http://networkx.github.io

12 http://ginsim.org/models_repository

13 The notebook can be previewed and downloaded at https://nbviewer.jupyter.org/github/colomoto/colomoto-docker/blob/2018-03-31/tutorials/CellCollective/CellCollective%20-%20Knowledge%20Base.ipynb

14 The notebook can be previewed and downloaded at https://nbviewer.jupyter.org/github/colomoto/colomoto-docker/blob/2018-03-31/tutorials/GINsim/GINsim%20-%20visualization.ipynb

15 The notebook can be previewed and downloaded at https://nbviewer.jupyter.org/github/colomoto/colomoto-docker/blob/2018-03-31/tutorials/bioLQM/bioLQM_tutorial.ipynb

16 The notebook can be previewed and downloaded at https://nbviewer.jupyter.org/github/colomoto/colomoto-docker/blob/2018-03-31/tutorials/Pint/quick-tutorial.ipynb

17 The notebook can be previewed and downloaded at https://nbviewer.jupyter.org/github/colomoto/colomoto-docker/blob/2018-03-31/tutorials/NuSMV/NuSMV%20with%20GINsim.ipynb

18 The notebook can be previewed and downloaded at https://nbviewer.jupyter.org/github/colomoto/colomoto-docker/blob/2018-03-31/tutorials/MaBoSS/MaBoSS%20-%20Quick%20tutorial.ipynb

19 The notebook can be previewed and downloaded at https://nbviewer.jupyter.org/github/colomoto/colomoto-docker/blob/2018-03-31/tutorials/GINsim/GINsim%20-%20visualization.ipynb

20 The notebook can be previewed and downloaded at https://nbviewer.jupyter.org/github/colomoto/colomoto-docker/blob/2018-03-31/tutorials/MaBoSS/Predict%20mutations%20with%20Pint,%20refine%20with%20MaBoSS.ipynb

21 http://jupyter.org

22 The notebook can be previewed and downloaded at https://nbviewer.jupyter.org/github/colomoto/colomoto-docker/blob/2018-03-31/tutorials/bioLQM/Fixpoints%20(interactive).ipynb

23 The notebook can be previewed and downloaded at https://nbviewer.jupyter.org/github/colomoto/colomoto-docker/blob/2018-03-31/tutorials/Reproducibility%20-%20fixpoints.ipynb

24 The notebook can be previewed and downloaded at https://nbviewer.jupyter.org/github/colomoto/colomoto-docker/blob/2018-03-31/tutorials/Reproducibility%20-%20model%20checking.ipynb

25 you may have to use pip3 instead of pip depending on your configuration

26 If using Docker Toolbox, the command should be executed within the Docker Terminal

27 https://gist.github.com

28 https://www.myexperiment.org

29 https://zenodo.org

30 Guidelines available at https://github.com/colomoto/colomoto-docker/blob/master/CONTRIBUTING.md

